# Temporal regulation of dUTP biosynthesis limits uracil incorporation during early DNA replication

**DOI:** 10.1101/027508

**Authors:** D. Suzi Bryan, Jay R. Hesselberth

## Abstract

Antimetabolite chemotherapies increase uracil levels in DNA, and thus identification of factors that influence the uracil content in DNA may have implications for understanding uracil-mediated chromosomal instability. We previously showed in the budding yeast *Saccharomyces cerevisiae* that uracil content in DNA correlates with replication timing, where the earliest and latest replicating regions are depleted in uracil. Here, we manipulated nucleotide biosynthesis enzymes in budding yeast to determine whether the pattern of uracil incorporation could be altered. In strains with high levels of uracil incorporation, deletion of dCMP deaminase (Dcd1) accelerated uracil incorporation at early-firing origins, likely due to rapid dTTP pool depletion. In contrast, increasing the activity of ribonucleotide reductase, which is required for the synthesis of all dNTPs via ribonucleotide diphosphates, lead to dUTP and dTTP pool equilibration and a concomitant increase in uracil content throughout the genome. These data suggest that uracil availability and the dUTP:dTTP ratio are temporally regulated during S phase and govern uracil incorporation into the genome. Therapeutic manipulation of nucleotide biosynthesis in human cells to either increase the dUTP pool or deplete the dTTP pool in early S phase may therefore improve the efficacy of antimetabolite chemotherapies.

## INTRODUCTION

To sustain DNA replication, cells must synthesize or scavenge precursors to accumulate intracellular pools of deoxynucleotide triphosphates (dNTPs). Nucleotide pools must be maintained above a threshold level during DNA synthesis (Koc et al. 2004), which requires multiple rounds of dNTP production in order to replicate the genome in its entirety (Poli et al. 2012). Deoxythymidine triphosphate (dTTP) is one of the four dNTPs essential for DNA replication. The critical step in the biosynthetic pathway of dTTP biosynthesis is the conversion of dUMP to dTMP, which is catalyzed by the enzyme thymidylate synthase (TS, **Figure 1**). Thymidylate synthase is a temporally regulated enzyme whose activity peaks in mid S phase of the cell cycle, and is required to sustain cell growth (Storms et al. 1984). In one branch of this pathway, sequential action by ribonucleotide reductase (RNR) and nucleoside diphosphate kinase (Ynk1) results in the biosynthesis of deoxyuridine triphosphate (dUTP), which is a natural precursor of dTTP and can be incorporated directly into DNA to create A:U base pairs, as the replicative polymerase does not discriminate between dUTP and dTTP (**Figure 1**) (Warner et al. 1981; Tinkelenberg et al. 2002; Lasken et al. 1996; Greagg et al. 1999; Wardle et al. 2008; Matuo et al. 2010).

**Figure 1.**
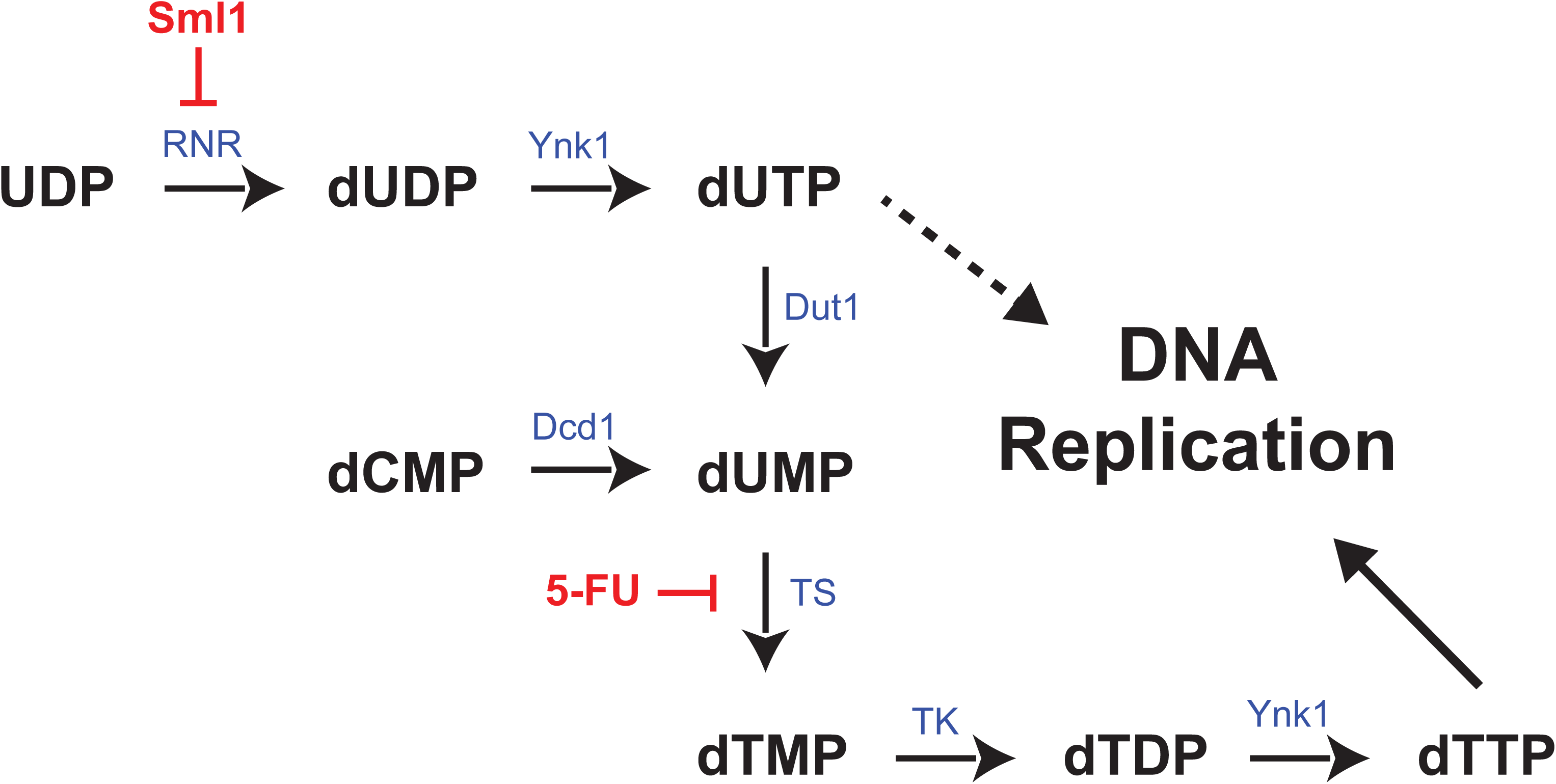
Biosynthesis of dUTP and dTTP. dTTP is one of the four canonical nucleobases used for DNA replication. Uracil in the form of dUTP is a natural precursor of dTTP that can be directly incorporated into the genome. Key enzymes in this pathway include ribonucleotide reductase (RNR), Nucleoside diphosphate kinase (Ynk1), dUTPase (Dut1), Deoxycitidylate deaminase (Dcd1), Thymidylate kinase (TK), and Thymidylate synthase (TS). RNR is inhibited by Sml1 in G_2_/M phase of the cell cycle. Antifolate chemotherapies like 5-fluorouracil (5-FU) covalently inhibit TS to induce dUTP accumulation, DNA damage and apoptosis.

In ribonucleotide reductase-mediated dTTP synthesis, production of dUMP is catalyzed by dUTP pyrophosphatase (Dut1). Dut1 is an essential enzyme necessary to maintain viability and normal nucleotide metabolism, and is expressed at a relatively constant level during the cell cycle (McIntosh et al. 1986; Guillet et al. 2006). In the absence of Dut1, cells accumulate dUTP and deplete dTTP pools (Gadsden et al. 1993), causing a futile cycle of dUTP incorporation and excision repair that leads to extensive DNA damage (Kavli et al. 2007). In S. *cerevisiae, dut1* null mutants cannot synthesize dTMP for the production of dTTP and are therefore only viable when supplemented with exogenous dTMP (Gadsden et al. 1993).

Independent of dUTP production, dUMP can be synthesized through deoxycytidylate (dCMP) deaminase (Dcd1). Dcd1 is an enzyme that produces dTTP throughout the cell cycle via conversion of dCMP to dUMP, and has the capability to produce adequate quantities of dTTP to sustain DNA replication (McIntosh and Haynes 1984). Unlike thymidylate synthase, dCMP deaminase is expressed at a constant level during the cell cycle, enabling Dcd1 to expand the dUMP pool both during and between successive S phases during cell division (**Figure 1**) (McIntosh and Haynes 1986; McIntosh et al. 1986).

Multiple nucleotide precursors feed into dUTP and dTTP biosynthesis pathways. It is also important to note that many enzymatic steps in dUTP and dTTP biosynthesis may operate bidirectionally; for example, dUTP is synthesized from dUMP using multiple intracellular kinases (Garces and Cleland 1969; Chien et al. 2009). Therefore, dUTP and dTTP pools are in a constant state of flux and are dependent on multiple levels of regulation.

When uracil is present in the genome, the repair enzyme uracil DNA glycosylase (UDG) recognizes and removes the uracil base, cleaving its glycosidic bond and leaving an abasic site (Krokan *et al.* 2002; Kavli *et al.* 2007). Inactivation of both uracil DNA glycosylase and dUTP pyrophosphatase in *E. coli* yields viable cells that stably incorporate uracil into their DNA, resulting from a buildup of dUTP and lack of a repair enzyme to recognize and remove uracil from DNA (Warner *et al.* 1981). Mutation of UDG and Dut1 have been shown to have similar effects on DNA of S. *cerevisiae* (Guillet *et al.* 2006) and *C. elegans* (Dengg *et al.* 2006). Increased dUTP incorporation is observed in regions of high transcription, suggesting that the levels of UTP necessary to sustain transcription may be converted to dUTP for incorporation into DNA (Kim and Jinks-Robertson 2009).

Pharmacologic treatments promote dUTP incorporation into DNA in order to cause DNA damage and cancer cell death. Antimetabolite drugs such as pemetrexed and 5-fluorouracil inhibit thymidylate synthase, increasing dUMP and dUTP levels (Longley *et al.* 2003). Because thymidylate synthase uses tetrahydrofolate as a methyl donor, folate deficiency also alters the pool of available nucleoside precursors for dTTP synthesis by increasing the cellular levels of dUTP, causing its incorporation into DNA and subsequent genome instability from upregulated uracil excision (Blount *et al.* 1997).

We previously developed Excision-seq, a new method for mapping DNA modifications that combines DNA treatment with excision-repair enzymes and high throughput DNA sequencing to identify content of modified nucleobases with high precision (Bryan *et al.* 2014). We applied this method to map uracil in genomic DNA and discovered a positional bias in genomic uracil content in a strain of yeast that accumulates an abundance of uracil in its genome. Here, we manipulated nucleotide biosynthetic pathways during the cell cycle to characterize the mechanism of variation in genomic uracil content.

## RESULTS

We previously used Excision-seq to map uracil in the yeast genome, and discovered a positional bias in genomic uracil content in a strain of yeast that accumulates an abundance of uracil in its genome: the earliest replicating regions exhibit a striking reduction in uracil content. In addition, the latest replicating regions in yeast also show a modest depletion of uracil (Bryan *et al.* 2014). Sequence read coverage in post-digestion Excision-seq data is inversely proportional to content of the modified nucleobase: regions with high levels of coverage have low levels of the modified base, and *vice versa.*

To determine whether temporal nucleotide biosynthesis was responsible for mediating genomic uracil incorporation, we deleted the key dNTP biosynthesis enzymes *DCD1* and/or *SML1* from *dut1-1 ung1*Δ yeast. *DCD1* encodes dCMP deaminase, which provides a pathway for dTTP biosynthesis that does not use dUTP as an intermediate (**Figure 1**). Because *DCD1* can produce adequate amounts of dTTP to sustain DNA replication (McIntosh and Haynes 1984), we predicted that removing this dTTP synthesis pathway would significantly reduce the dTTP pool and increase the dUTP:dTTP ratio in early S phase, leading to an increase uracil incorporation at replication origins. *SML1* is an inhibitor of ribonucleotide reductase (RNR), a temporally regulated enzyme necessary for synthesizing all dNTPs used during DNA replication (**Figure 1**) (Zhao et al. 1998; Chabes et al. 1999; Zhao et al. 2000). We anticipated that increasing the level of all intracellular dNTP pools will drive the dUTP:dTTP ratio toward equilibrium, resulting in less uracil content variation genome-wide.

Specifically, we anticipated uracil incorporation at origins of replication and late-replicating regions in *dut1-1 ung1*Δ *sml1*Δ.

### Mutations in nucleotide biosynthesis enzymes affect cell viability

To determine whether modifying nucleotide biosynthesis via deletion of dCMP deaminase (*DCD1*) or deletion of the ribonucleotide reductase inhibitor *SML1* affected normal cellular function and DNA replication, we performed a dilution analysis growth assay in mutants of the biosynthetic pathway grown on YEPD and 50, 100, and 200 mM hydroxyurea. Hydroxyurea (HU) is an inhibitor of ribonucleotide reductase that prevents dNTP pool expansion in G_1_/S that is necessary to sustain DNA replication (Koc et al. 2004). When nucleotide biosynthesis mutants were exposed to HU, cellular viability was altered. Growth of wild type yeast was arrested on HU because nucleotide pools were depleted below the threshold level required for DNA replication (**Figure 2A**) (Koc et al. 2004). Deletion of *SML1* rescued growth, as expected from previous studies (Zhao *et al.* 1998) because upregulation of ribonucleotide reductase increases nucleotide pool levels above the threshold level necessary to maintain DNA replication (Koc et al. 2004) (**Figure 2A**). However, deletion of *SML1* in *dut1-1 ung1*Δ yeast did not rescue growth, and had similar growth to wild type cells on HU (**Figure 2A**). The lack of rescued growth in *dut1-1 ung1*Δ *sml1*Δ yeast was surprising, and potentially indicates a requirement for thymine in cells that can stably incorporate uracil into DNA.

**Figure 2.**
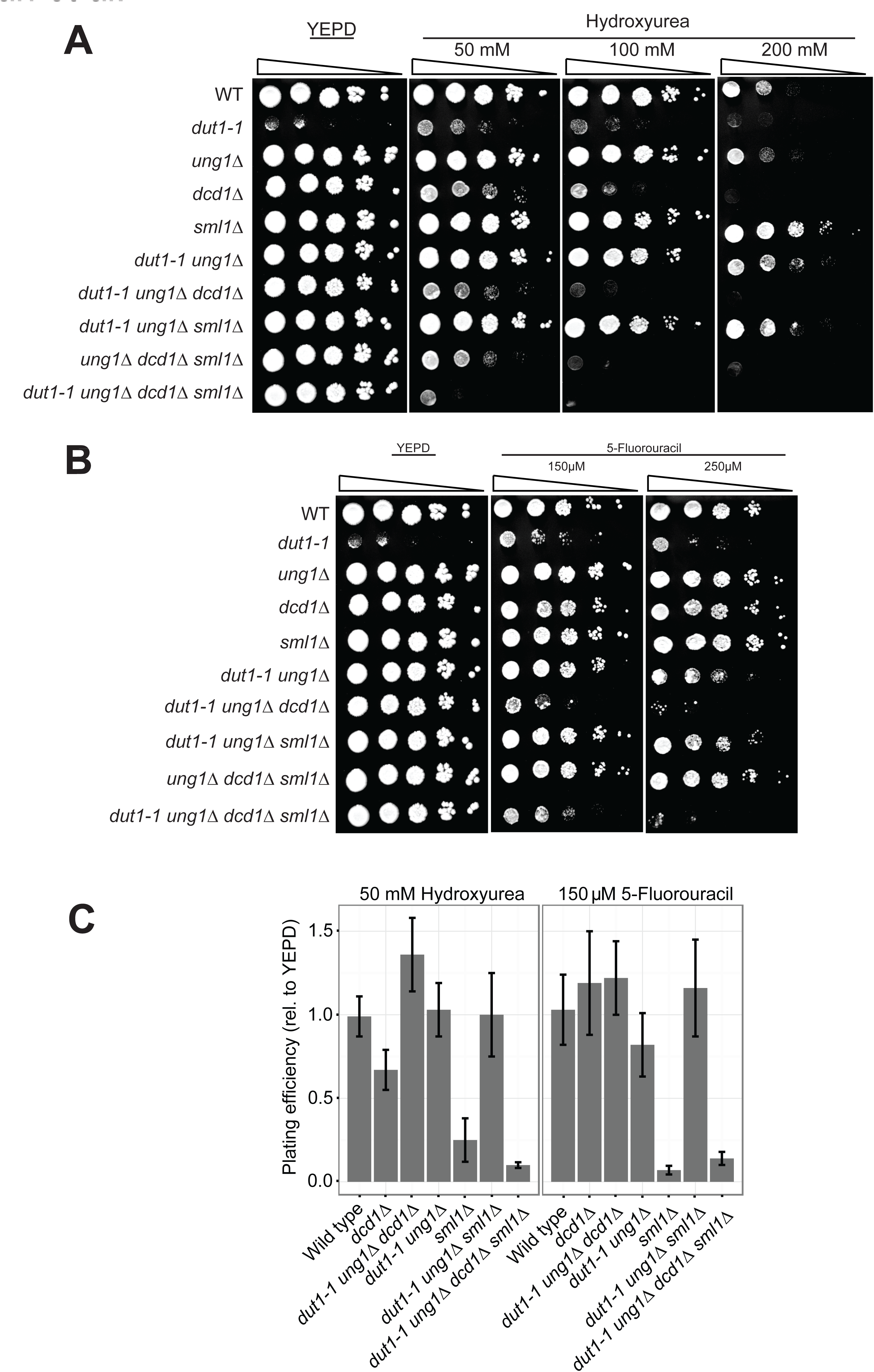
Growth assays for dNTP biosynthesis mutants. **A.** Yeast strains were plated on YEPD and hydroxyurea and grown for three days at 30°C before imaging. Five 1:10 serial dilutions of wild type, single, double, triple, and quadruple mutant strains are shown. **B.** Yeast strains were plated on YPD and 5-fluorouracil and grown for three days at 30°C before imaging. Five 1:10 serial dilutions of wild type, single, double, triple, and quadruple mutant strains are shown. **C.** Quantitative efficiency of plating assay shows percent viability of single, double, and triple mutant strains relative to wild type on 50 mM hydroxyurea (HU) and 150 μM 5-fluorouracil (5-FU).

In order to determine the individual contributions of each nucleotide biosynthesis enzyme to cellular viability, we performed an efficiency of plating assay for wild type and single, double, triple and quadruple mutants of *DCD1* and *SML1* in *dut1-1, ung1*Δ yeast. This assay demonstrated that *sml1*Δ yeast plating efficiency increased 1.4-fold on 50 mM HU relative to YEPD, while wild type efficiency did not change (**Figure 2C**). In contrast, *dut1-1 ung1*Δ *sml1*Δ yeast showed equivalent plating efficiency compared with wild type when plated on 50 mM HU (**Figure 2C**), indicating that increased nucleotide production does not rescue growth in *dut1-1 ung1*Δ yeast under HU replicative stress, as demonstrated in qualitative assays. Therefore, these data reinforce the apparent requirement for dTTP in *sml1*Δ-mediated rescue of growth on hydroxyurea that was observed in qualitative cell viability assays (**Figure 2C**).

*DCD1* is necessary for dUTP-independent synthesis of dTTP, and produces sufficient dTTP to maintain DNA replication (McIntosh and Haynes 1984). We expected that treatment of yeast that lack this branch of dTTP biosynthesis with hydroxyurea would exacerbate the phenotypic effects of *DCD1* deletion; i.e., viability would be reduced. Single mutant *dcd1*Δ yeast and triple mutant *dut1-1 ung1*Δ *dcd1*Δ yeast showed increased sensitivity to HU treatment, likely resulting from depletion of their dTTP pool sooner than wild type and *dut1-1 ung1*Δ yeast (**Figure 2A**). A quantitative plating assay for single mutant *dcd1*Δ yeast showed a 1.5-fold decrease in plating efficiency relative to YEPD on 50 mM HU, while triple mutant *dut1-1 ung1*Δ *dcd1*Δ decreased to only 0.25-fold plating efficiency relative to YEPD on HU (**Figure 2C**). These data support our hypothesis that removal of *DCD1*-mediated production of dTTP increases the intracellular dUTP:dTTP ratio in early S phase, and is consistent with cells having a requirement for dTTP even when dUTP can be stably substituted in the genome.

To evaluate the combined influences of *DCD1* and *SML1* on strain viability, we deleted both enzymes to create a quadruple mutant *dut1-1 ung1*Δ *dcd1*Δ *sml1*Δ strain. The mutant *dut1-1 ung1*Δ *dcd1*Δ *sml1*Δ strain exhibited high HU sensitivity, showing a greater growth defect than even *dut1-1 ung1*Δ *dcd1*Δ yeast (**Figure 2A**). Quantitatively, *dut1-1 ung1*Δ *dcd1*Δ *sml1*Δ yeast plating efficiency was decreased to only 0.10-fold relative to YEPD on HU, a 2.5-fold decrease in viability compared to *dut1-1 ung1*Δ *dcd1*Δ yeast (**Figure 2C**). Since deletion of *SML1* should decrease HU sensitivity, this result was unexpected, and is possibly a result of the decreased dTTP pool in G_1_ via removal of *DCD1,* whose protein product is normally constitutively active during the cell cycle and able to create dTTP (via dUMP) in sufficient quantities to sustain replication (McIntosh and Haynes 1986; McIntosh et al. 1986; McIntosh and Haynes 1984; Kudlicki et al. 2007). Triple mutant *ung1*Δ *dcd1*Δ *sml1*Δ yeast also showed moderate sensitivity to HU, further demonstrating the importance of Dcd1 in dTTP production, even when the pool of dUTP is not increased from the *dut1-1* mutation (**Figure 2B**).

Overall reduction of dNTPs for DNA replication via hydroxyurea demonstrated a surprising dTTP requirement in uracil-rich yeast. To further characterize the necessity of dTTP in *dut1-1 ung1*Δ yeast, we aimed to specifically decrease the dTTP pool by growing yeast strains in the presence of 150 and 250 μM 5-fluorouracil (5-FU), an antifolate drug used in chemotherapy to reduce dTTP production and therefore increase cellular levels of dUTP to induce chromosomal instability (Blount et al. 1997). This treatment mirrors uracil accumulation in a *dut1-1 ung1*Δ strain, but causes genome instability in strains with wild type Ung1 (Seiple et al. 2006). We predicted that the requirement for dTTP would be further emphasized in uracil-abundant strains by exposing them to 5-FU. When a *dut1-1 ung1*Δ strain was treated with 5-fluorouracil, cell viability decreased relative to wild type (**Figure 2B**). This result was again surprising since dUTP should invisibly substitute for dTTP in this strain, and supports the cellular requirement for dTTP that we previously proposed. A quantitative plating assay showed that *dut1-1 ung1*Δ yeast plating efficiency was 0.82-fold relative to YEPD on 150 μM 5-FU, whereas wild type efficiency showed no change (**Figure 2C**). This sensitivity phenotype was rescued by deletion of *SML1* in *dut1-1 ung1*Δ yeast, which showed a 1.2-fold increase in plating efficiency relative to YEPD on 150 μM 5-FU (**Figure 2B** and **Figure 2C**). Because *SML1* deletion and subsequent RNR upregulation expands all dNTP pools, we expected growth to be rescued in *dut1-1 ung1*Δ *sml1*Δ yeast, indicating that the *sml1*Δ-mediated expansion of the intracellular dTTP pool restored the dTTP level above the necessary threshold to sustain DNA replication (Koc et al. 2004).

As discussed, dCMP deaminase (*DCD1*) produces dTTP to sustain DNA replication (McIntosh and Haynes 1984); therefore, we hypothesized that deletion of this dTTP biosynthesis pathway would have significant detrimental consequences when uracil-abundant strains were exposed to pharmacological dTTP depletion via 5-FU. Single mutant *dcd1*Δ yeast did not show a qualitative or quantitative decrease in viability relative to YEPD on 5-FU, indicating that dTTP is produced in sufficient amounts through RNR and dUTPase in yeast with wild type *DUT1* and *UNG1* to withstand moderate pharmacological dTTP depletion (**Figure 2B** and **Figure 2C**). Unlike *dcd1*Δ yeast, triple mutant *dut1-1 ung1*Δ *dcd1*Δ yeast showed increased sensitivity to 5-FU compared with *dut1-1 ung1*Δ and wild type yeast, potentially resulting from the combined effects of depleting dTTP in G_1_ via *DCD1* deletion and depleting the dUMP pool for dTTP production via *DUT1* modification (**Figure 2B**) Quantitatively, *dut1-1 ung1*Δ *dcd1*Δ yeast had only 0.072-fold plating efficiency relative to YEPD on 5-FU (**Figure 2C**). These data suggest that the effect of deleting dCMP deaminase in the presence of a dTTP biosynthesis inhibitor like 5-FU on cellular viability is significantly intensified in yeast with increased genomic uracil.

To determine whether the sensitivity of *dut1-1 ung1*Δ *dcd1*Δ yeast on 5-FU could be rescued by increasing the dTTP pool, we exposed quadruple mutant *dut1-1 ung1*Δ *dcd1*Δ *sml1*Δ yeast to 5-FU. While *dut1-1 ung1*Δ *dcd1*Δ *sml1*Δ yeast exhibited 5-FU sensitivity, it was moderately less than the *dut1-1 ung1*Δ *dcd1*Δ mutant, indicating that the increase in the dTTP pool by RNR upregulation and subsequent dTTP incorporation in *dut1-1 ung1*Δ *dcd1*Δ yeast increases cellular viability (**Figure 2B**). Quantitatively, *dut1-1 ung1*Δ *dcd1*Δ *sml1*Δ yeast showed a plating efficiency of 0.14-fold relative to YEPD on 5-FU, a 2-fold increase in viability compared to *dut1-1 ung1*Δ *dcd1*Δ yeast (**Figure 2C**). The moderate rescue phenotype in *dut1-1 ung1*Δ *dcd1*Δ *sml1*Δ yeast further reinforces the requirement for dTTP in uracil-rich yeast genomes. In addition, it emphasizes that the necessity for dTTP is highest in early S phase, correlating with the transition from normal to slow replicating mode observed in yeast exposed to hydroxyurea (Poli et al. 2012). Since the activation of 5-FU to FdUMP requires ribonucleotide reductase, dTTP depletion via thymidylate synthase inhibition does not occur until after RNR activation in S phase. Increased dTTP availability when DNA replication begins results in the rescue in viability observed when *SML1* was deleted from *dut1-1 ung1*Δ *dcd1*Δ yeast.

Overall, these data indicate that cells require a threshold level of dTTP to maintain proper cellular functions, even when dUTP can be substituted stably in the DNA of yeast lacking the uracil repair enzyme Ung1. The surprising sensitivity of the *dut1-1 ung1*Δ *dcd1*Δ *sml1*Δ mutant on both HU and 5-FU indicates that the temporal regulation of dNTP biosynthesis pathways influences cell viability and likely DNA composition based on dNTP levels available for nascent DNA synthesis at a given time, most significantly in G_1_ and early S phase of the cell cycle.

### Excision-seq mapping of nucleotide biosynthesis mutants

During the cell cycle, nucleotide biosynthesis is temporally regulated by multiple enzymes in order to maintain dNTP pools (Storms et al. 1984; McIntosh et al. 1986). We hypothesized that variation in genomic uracil content revealed by Excision-seq mapping is the result of changes in dUTP and dTTP pool availability during G_1_ and S phase. In order to determine whether uracil content was altered when nucleotide pools changed, we performed post-digestion Excision-seq on *dut1-1 ung1*Δ *dcd1*Δ yeast (S288C background) and *dut1-1 ung1*Δ *sml1*Δ yeast (S288C and W303 backgrounds) and mapped reads to the yeast genome for visualization in the UCSC genome browser (Karolchik *et al.* 2001). We mapped normalized uracil depletion signal (reads per million, RPM) and compared the data with that of *dut1-1 ung1*Δ yeast. Using post-digestion Excision-seq, read coverage is inversely proportional to uracil content; therefore, regions with high levels of sequencing reads indicate low levels of uracil, and *vice versa.* We examined alterations to genomic uracil content when regulatory dNTP biosynthesis enzymes were altered or removed. When *DCD1* was deleted, uracil was incorporated into DNA sooner after replication origins fire, indicative of the reduced dTTP pool and increased dUTP:dTTP ratio in G_1_ (**Figure 3A**, green). Deletion of *SML1* results in increased ribonucleotide reductase (RNR) activity because it no longer binds to and inhibits RNR in G_2_/M phase of the cell cycle. Removing Sml1 inhibition from RNR abolishes the rate-limiting step for dNTP biosynthesis, resulting in an overall increase in total dNTP levels (Zhao et al. 2001). When Excision-seq uracil mapping was performed in *dut1-1 ung1*Δ *sml1*Δ yeast DNA, we observed a more distributed pattern of uracil incorporation throughout the genome, with higher uracil levels at origins and slightly lower uracil levels in areas surrounding origins (**Figure 3A**, blue and orange).

**Figure 3.**
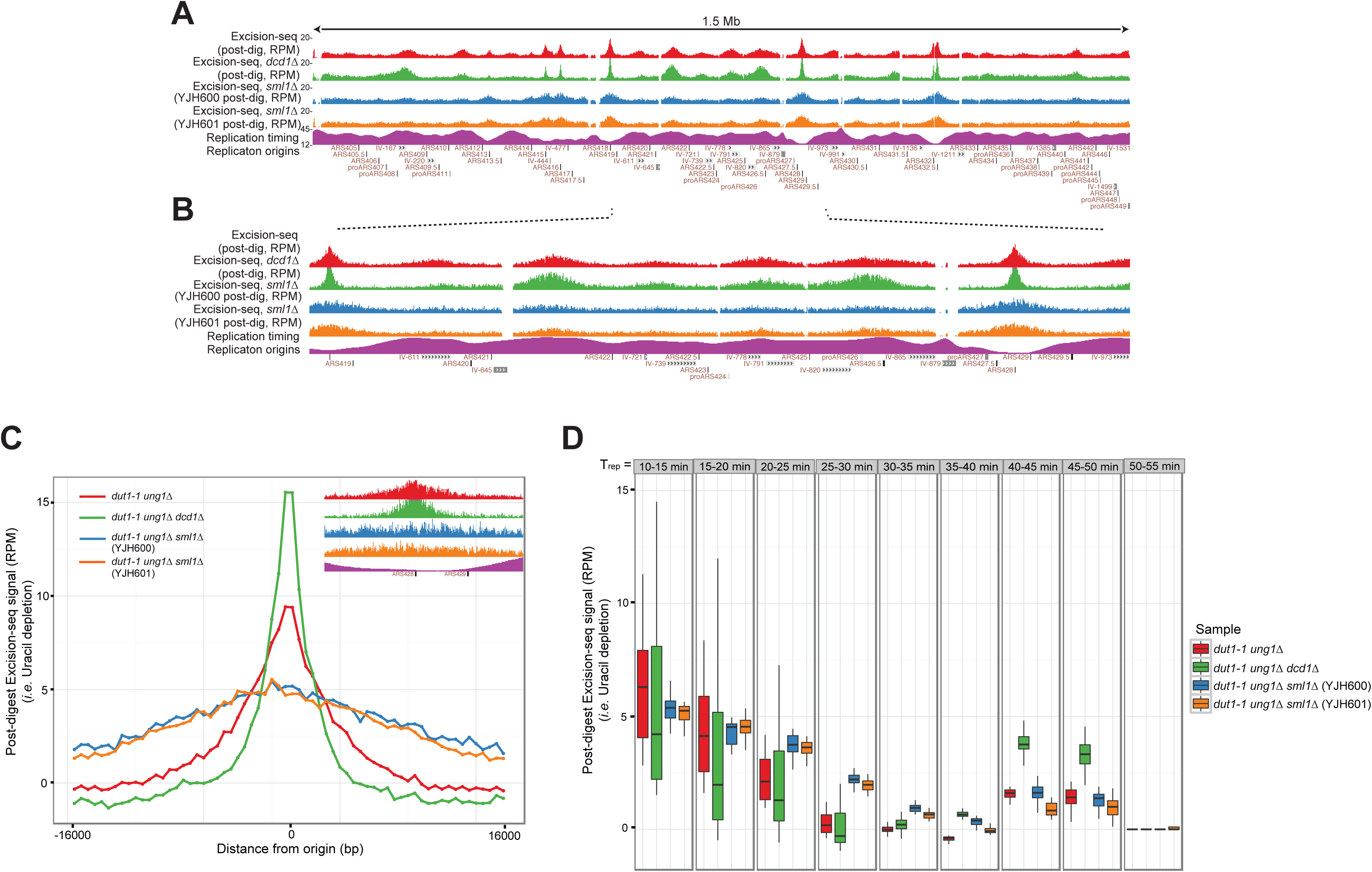
Temporal regulation of nucleotide biosynthesis via *DCD1* and *SML1*. **A.** Data showing the entire yeast chromosome 4 using post-digestion Excision-seq for *dut1-1 ung1*Δ yeast (red, reads per million (RPM)), *dut1-1 ung1*Δ *dcd1*Δ yeast (green, RPM), *dut1-1 ung1*Δ *sml1*Δ yeast (blue and orange, RPM), replication timing data (T_rep_, minutes replicated after G_1_ release) (Raghuraman 2001) (grey), and annotated origins of replication (Nieduszynski et al. 2007). **B.** A 450 kb region of chromosome 4 highlights patterns of uracil incorporation in early replicating origins (*ARS418* and *ARS428*), as well as uracil depletion in late replicating regions. **C.** Quantitative meta-analysis of a ~32 kb region showing the average Excisionseq (uracil depletion, RPM) signal for 42 early firing origins of replication. Insert: 32kb surrounding representative origin ARS428. Red: *dut1-1 ung1*Δ, green: *dut1-1 ung1*Δ *dcd1*Δ, blue: *dut1-1 ung1*Δ *sml1*Δ (S288C background), orange: *dut1-1 ung1*Δ *sml1*Δ (W303 background), pink: replication timing data (T_rep_, minutes replicated after G_1_ release) (Raghuraman 2001), and annotated origins of replication (Nieduszynski et al. 2007). **D.** Genome-wide meta-analysis separated into 5 minute replication timing windows of ~10 kb regions. Red: *dut1-1 ung1*Δ, green: *dut1-1 ung1*Δ *dcd1*Δ, blue: *dut1-1 ung1*Δ *sml1*Δ (S288C background), orange: *dut1-1 ung1*Δ *sml1*Δ (W303 background).

To determine whether uracil depleted regions changed genome-wide when dNTP biosynthesis was altered, we compared the ratio of Excision-seq signal (*i.e.* uracil depletion) for early-firing origins compared with mid-replicating DNA ~10 kb away, an indicator of uracil depletion levels at replication origins and uracil enrichment levels at mid-replicating genomic regions. When *DCD1* was deleted from *dut1-1 ung1*Δ yeast, this ratio increased ~4-fold (**Figure 3A**, **B**, green, ARS428). Although unexpected, this result is explained by the bidirectionality of the dUTP to dUMP reaction (**Figure 1**) (Garces and Cleland 1969; Chien et al. 2009). Deletion of Dcd1 results in less dUMP in G_1_ and therefore less flux from dUMP to dUTP in this pathway, resulting in a higher peak of uracil depletion at replication origins (**Figure 3A**, **B**, green). In *dut1-1 ung1*Δ *sml1*Δ yeast, the ratio of early to mid-replicating DNA decreased ~4-fold compared with *dut1-1 ung1*Δ yeast (**Figure 3A, B,** blue and orange, ARS428). This result corroborates the equilibration of dTTP and dUTP, resulting in more uracil at origins (*i.e.* less sequencing reads) and marginally less uracil in mid-replicating genomic regions (**Figure 3A, B,** blue and orange). A comparison of late-replicating regions to nearby mid-replicating regions showed that deletion of *DCD1* or *SML1* in *dut1-1 ung1*Δ yeast did not significantly change the ratio of late to mid-replicating DNA (**Figure 3A**, green, blue and orange, peaks between ARS406 and 409).

### Quantification of uracil depletion in nucleotide biosynthesis mutants

Excision-seq mapping of *dut1-1 ung1*Δ yeast revealed 42 early-firing origins depleted of uracil across the genome (Bryan *et al.* 2014). In order to determine average uracil depletion for all uracil depleted origins genome-wide, we performed a metadata analysis to examine the average post-digestion Excision-seq signal for *dut1-1 ung1*Δ, and compared these data to *dut1-1 ung1*Δ *dcd1*Δ and *dut1-1 ung1*Δ *sml1*Δ yeast across a ~32 kb genomic region. This region encompassed uracil depletion peaks for all dNTP biosynthesis mutants (**Figure 3C**). Within this region, we calculated peak thresholds to identify uracil depletion peaks for *dut1-1 ung1*Δ*, dut1-1 ung1*Δ *dcd1*Δ, and *dut1-1 ung1*Δ *sml1*Δ yeast strains relative to background levels (*i.e.* low signals in mid-replicating, uracil-rich regions).

*DCD1* is responsible for maintaining the dTTP pool in G_1_ and early S phase. We predicted that removing this enzyme would significantly reduce dTTP pools during this time, resulting in uracil incorporation sooner after replication began. We analyzed uracil incorporation near origins by *DCD1* deletion, and determined that uracil depletion peak width decreased from ~6 kb for *dut1-1 ung1*Δ yeast to ~2.5 kb for *dut1-1 ung1*Δ *dcd1*Δ yeast (**Figure 3C,** green). We also calculated the rate of uracil incorporation after replication origins fire, and determined that the incorporation rate for the *dcd1*Δ mutant strain increased by an average of ~3-fold when compared with that of *dut1-1 ung1*Δ yeast (**Figure 3C,** green). These data indicate that dTTP is being depleted faster and therefore dUTP is being incorporated sooner after DNA replication is initiated. Directly at origins, yeast deleted for *DCD1* show an increase in average sequencing signal relative to *dut1-1 ung1*Δ yeast (*i.e.* greater uracil depletion). Once again, this result is explained by the lack of dCMP deaminase in G_1_ and the bidirectional potential of nucleotide biosynthesis pathways. Because *DCD1* is not deaminating dCMP to dUMP, less dUTP is being generated from dUMP before and during initial DNA replication (**Figure 1**).

Removal of *SML1* from yeast strains increases ribonucleotide reductase activity during the cell cycle and increases the availability of all dNTPs. We hypothesized that deletion of *SML1* would drive the dUTP:dTTP ratio toward equilibrium, and uracil would be incorporated at replication origins. Deletion of *SML1* in *dut1-1 ung1*Δ yeast resulted in significantly lower average signal at replication origins (*i.e.* higher in uracil content), indicating that uracil and thymine are more distributed throughout the genome through dNTP pool equilibration (**Figure 3C,** blue and orange). Quantitative analysis showed that the rate of uracil incorporation for the *SML1* deleted strains decreased by an average of ~7-fold when compared with that of *dut1-1 ung1*Δ yeast (**Figure 3C,** blue and orange). These data indicate that dUTP is not significantly depleted at origins when ribonucleotide reductase is upregulated via *SML1* deletion, and therefore dTTP levels are not diminished at an appreciable rate because the dUTP pool is not limiting when DNA replication is initiated.

Excision-seq mapping of uracil in *dut1-1 ung1*Δ yeast revealed uracil depletion most significantly at origins, but also modestly at late-replicating genomic regions (Bryan *et al.* 2014). Therefore, we wanted to examine the average post-digestion Excision-seq signal (*i.e.* uracil depletion) relative to replication timing across the entire genome. We hypothesize that uracil in late-replicating regions will not be significantly altered because the impact of *DCD1* and *SML1* deletion will be muted by multiple rounds of dNTP biosynthesis during S phase. Analysis of the average post-digestion Excision-seq signal separated by replication timing was performed genome-wide in *dut1-1 ung1*Δ*, dut1-1 ung1*Δ *dcd1*Δ, and *dut1-1 ung1*Δ *sml1*Δ strains (**Figure 3D**). The average Excision-seq signals for ~10 kb surrounding replication origins were separated into 5 minute replication timing windows across the entire genome. This metadata analysis revealed that deleting *DCD1* resulted on average in a ~2-fold increase in sequencing reads (*i.e.* decrease in uracil incorporation) at late-replicating regions (T_rep_ > 40 min, **Figure 3D**). Similar to the increase in height of uracil depletion peaks (*i.e.* decrease in uracil) directly at replication origins, this result can be explained by the reduction in the portion of the dUMP pool that can be phosphorylated to dUTP during DNA replication (**Figure 1**). Deleting *DCD1* did not change uracil content in mid-replicating regions (T_rep_ = 25-40 min), which may be explained by the expansion of dNTP pools masking the minimal decrease in the dUMP pool that results from *DCD1* deletion (**Figure 3D**).

Unlike the significant changes in uracil incorporation at origins (**Figure 3C**) and the modest changes in late-replicating regions in *dut1-1 ung1*Δ *dcd1*Δ yeast, deleting *SML1* did not result in a significant alteration to uracil content in late-replicating genomic regions (T_rep_ > 40 min, **Figure 3D**). However, in mid-replicating regions (T_rep_ = 25-40 min), uracil content decreased an average of ~3-fold in *dut1-1 ung1*Δ *sml1*Δ strains. We postulate that this decrease is the result of dUTP and dTTP pool equilibration, causing increased uracil incorporation at replication origins and decreased uracil content in mid-replicating regions (**Figure 3D**). If this theory is correct, it may explain the moderate uracil depletion that we observed in late-replicating regions of *dut1-1 ung1*Δ yeast (Bryan *et al.* 2014), which we propose is the result of dNTP pools shifting toward equilibrium through multiple rounds of dNTP synthesis. Therefore, we would not expect uracil content in late-replicating regions to be altered when *SML1* is deleted from *dut1-1 ung1*Δ yeast because dUTP and dTTP pool equilibration previously occurred in mid-replicating regions (**Figure 3A,B,** blue and orange).

Data obtained from analysis of the average Excision-seq signal genome-wide signify that deleting *DCD1* or *SML1* in *dut1-1 ung1*Δ yeast significantly changes uracil incorporation at replication origins and mid-replicating regions, but does not abolish uracil depletion in late-replicating regions. This result was anticipated because dUTP and dTTP pools have been replenished by additional dNTP synthesis throughout S phase, resulting in an increase in the availability of both nucleotides for DNA replication. However, deletion of *DCD1* resulted in a slight decrease in the uracil content of late regions, which is likely the consequence of bidirectional dNTP biosynthesis pathways and the reduction in the portion of the dUMP pool which would normally be created from dCMP during DNA replication (**Figure 1**).

Overall, these data show that changing cell cycle regulation of nucleotide biosynthesis alters uracil incorporation genome-wide. Deletion of enzymes that govern dUTP and dTTP biosynthesis, especially when DNA replication initiates, has profound effects on variation in genomic uracil content. Additional Excision-seq runs and subsequent metadata analyses were repeated for *dut1-1 ung1*Δ strains deleted for *DCD1* or *SML1,* and comparable results were obtained (data not shown). The ability to obtain equivalent results reinforces the validity of the Excision-seq method, and demonstrates that it is a reproducible, accurate measurement of genomic uracil.

## DISCUSSION

We previously developed a method to map DNA modifications genome wide using excision repair enzymes in order to analyze their specific impact on genomic context (Bryan *et al.* 2014). Application of the Excision-seq method to map uracil in DNA revealed a previously unknown variation in uracil content genome-wide. We postulated that the lack of uracil at replication origins and late-replicating regions was the result of temporally regulated biosynthesis of dNTPs used for DNA replication. In this study, the deletion of a regulatory enzyme and an enzyme inhibitor instrumental in the biosynthesis of dUTP and dTTP revealed changes in the uracil depletion pattern of the yeast genome, correlating with alterations to the dUTP pool and dUTP:dTTP ratio. Based on these data, we propose a model in which temporal regulation of nucleotide biosynthesis is responsible for the variation in genomic uracil content, and alteration of dNTP pools in G_1_/S phase changes the uracil pattern we observe using Excision-seq.

### Regulation of nucleotide biosynthesis

During G_1_/S phase of the cell cycle, nucleotide pools are compartmentalized in the nucleus and are limiting for DNA synthesis, allowing ~5 kb of DNA to be replicated at a time (Poli *et al.* 2012). Nucleotides must undergo additional rounds of synthesis throughout S phase to maintain dNTP levels sufficient to continue replication (Koc et al. 2004). Therefore, the composition of nucleotides available for incorporation into DNA is constantly changing during G_1_ and S phase. We have demonstrated via altering regulation of the dNTP biosynthesis enzymes Dcd1 and Sml1 that in *dut1-1 ung1*Δ yeast, the length of nascent DNA that is protected from uracil incorporation after origins fire can be reduced or essentially abolished, resulting from increasing the dUTP:dTTP ratio and subsequent uracil incorporation by a replicative polymerase that does not discriminate between the two pools (**Figure 3C**) (Lasken et al. 1996; Greagg et al. 1999; Wardle et al. 2008; Matuo et al. 2010). These data support our model in which nucleotide pools are compositionally pure of dUTP in G_1_, and accumulation of dUTP only occurs after activation of ribonucleotide reductase and resynthesis of nucleotides in S phase (**Figure 4**). Surprisingly, uracil was reduced directly at replication origins as well as in late-replicating regions when *DCD1* was deleted from *dut1-1 ung1*Δ yeast. Although these results appear counterintuitive, they may be explained by the decrease in intracellular dUMP when *DCD1* is deleted, because nucleotide biosynthesis pathways operate bidirectionally (Garces and Cleland 1969; Chien et al. 2009). Therefore, dUMP may be phosphorylated to dUTP using endogenous monophosphate and diphosphate kinases, and thus supplement the available pool of dUTP (**Figure 1**). *SML1* deletion and the ensuing upregulation in ribonucleotide reductase activity resulted in increased uracil incorporation at replication origins and decreased uracil content in mid-replicating regions. Importantly, removal of this enzyme did not change uracil content in late-replicating genomic regions (**Figure 3D**). We previously hypothesized that modest uracil depletion in late-replicating regions of *dut1-1 ung1*Δ yeast (Bryan *et al.* 2014) was the result of dUTP and dTTP pool equilibration from multiple rounds of dNTP synthesis. In *dut1-1 ung1*Δ *sml1*Δ yeast, dUTP and dTTP pool equilibration already occurs in mid-replicating regions, and therefore uracil incorporation into late-replicating regions is not altered when *SML1* is deleted from *dut1-1 ung1*Δ yeast (**Figure 3D**), which supports our equilibration hypothesis for *dut1-1 ung1*Δ yeast.

**Figure 4.**
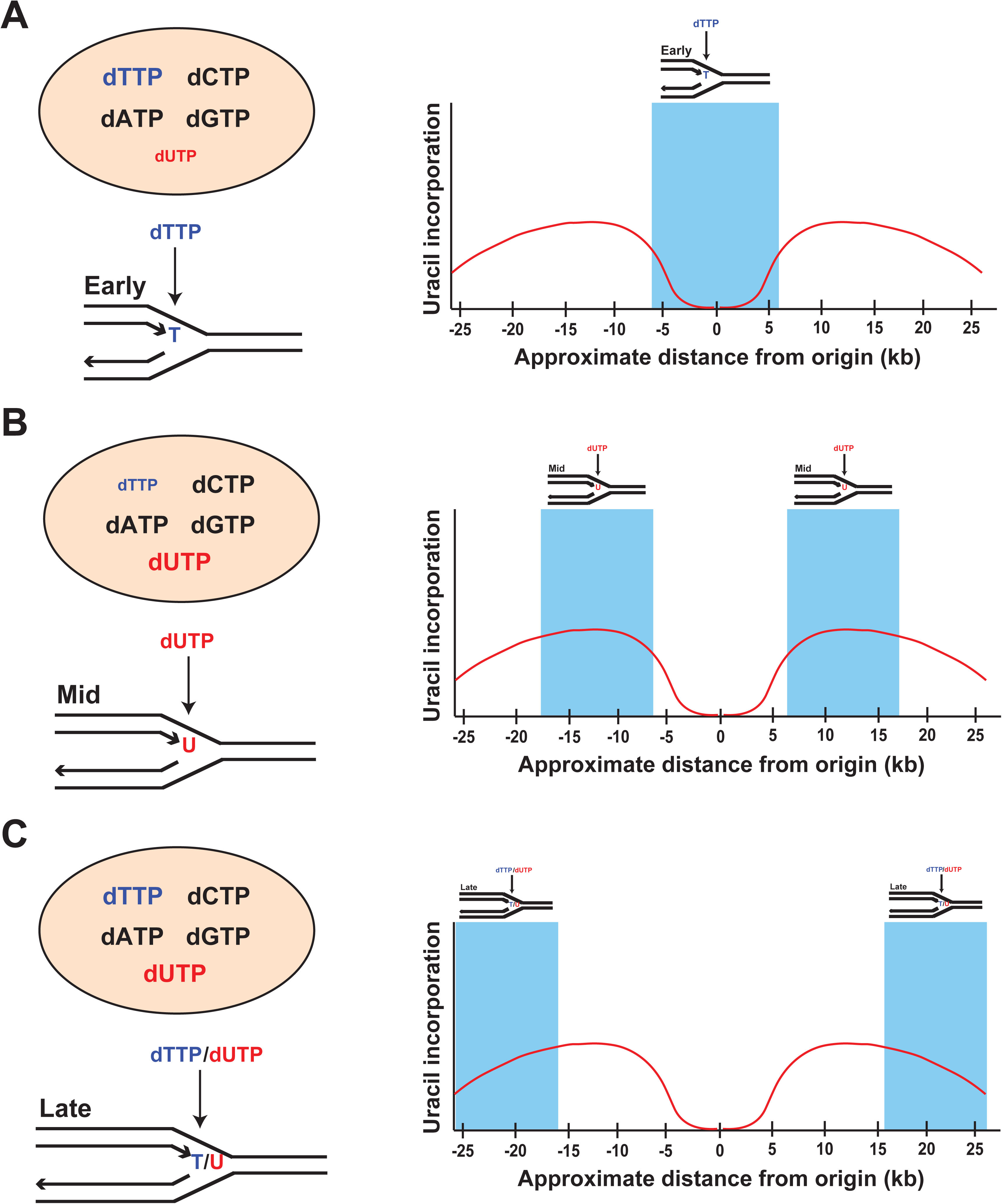
Model: nucleotide biosynthesis governs genomic uracil incorporation. **A.** At early-replicating genomic regions (*i.e.* origins of replication, shaded blue rectangle), the intracellular dNTP pool is compositionally pure of dUTP (red), and dTTP (blue) is abundant because of Dcd1 synthesis of dTMP. Therefore, dTTP is incorporated into the nascent DNA strand during replication. **B.** In mid-replicating regions (shaded blue rectangles), additional dNTP synthesis occurs via RNR upregulation. In a *dut1-1 ung1*Δ strain or an *ung1*Δ strain + 5-luorouracil, this equates to substantial dUTP (red) synthesis and incorporation into DNA. **C.** In late-replicating regions (shaded blue rectangles), dTTP (blue) and dUTP (red) pools partially equilibrate through multiple rounds of dNTP synthesis, resulting in competition with a nondiscriminatory polymerase for which dNTP is used during DNA replication.

The importance of temporal dTTP biosynthesis from both ribonucleotide reductase and dCMP deaminase is highlighted when dTTP pools are decreased. In wild type yeast, dCMP deaminase can supply adequate quantities of dUMP (for conversion to dTTP) to sustain cellular growth and viability (McIntosh and Haynes 1984). Our data from the growth analysis of mutant strains on hydroxyurea and 5-fluorouracil demonstrate that even in *dut1-1 ung1*Δ yeast, which stably incorporates uracil in place of thymine, a threshold level of dTTP is necessary to maintain normal cellular growth and function (**Figure 2**). Although this result is unexpected, cellular thymine requirement could possibly arise from alterations to cell cycle regulatory mRNA and protein gene products in uracil-rich genomes. Previous studies have demonstrated that when uracil is substituted for thymine in the genome, some genes lose the ability to efficiently bind transcription factors and other DNA binding proteins, leading to alterations in gene transcription levels (Gadsden *et al.* 1993). Therefore, one hypothesis is that the high level of uracil substitution for thymine present in *dut1-1 ung1*Δ yeast causes changes in the transcription of mRNA responsible for maintaining key cellular functions, especially under replication stress (*i.e.* hydroxyurea or 5-fluorouracil). In the future, a genome-wide analysis of gene transcription and/or protein expression may consequently reveal significant changes to cellular regulation in *dut1-1 ung1*Δ yeast.

### Implications for improvements to standard antimetabolite chemotherapy

The Excision-seq method of uracil mapping revealed variation in genomic uracil content that excludes large chromosomal regions near origins of replication from uracil incorporation, a phenomenon conserved in bacteria and yeast (Bryan *et al.* 2014). Preservation of uracil “cold spots” – sites of reduced uracil incorporation – in prokaryotic and eukaryotic organisms suggests that a similar uracil depletion pattern may be revealed via Excision-seq mapping in human cells treated with antimetabolite chemotherapies such as 5-fluorouracil, whose aim is to elevate cellular dUTP levels in order to promote excision by Ung1, DNA lesions, and ultimately apoptosis in cancer cells (Longley et al. 2003). Therefore, the ability to increase uracil levels at uracil “cold spots” in human cells using enzymes that alter dNTP biosynthesis to favor dUTP over dTTP incorporation may significantly improve the efficacy of chemotherapeutics that cause genomic uracil integration, chromosomal instability, and cell death.

## MATERIALS AND METHODS

### Yeast growth assays

*S. cerevisiae* liquid cultures (YJH259, 749, 506, 619, 755, 532, 599, 600, 756, and 752, **Table 1**) were grown in YEPD overnight and diluted in sterile 96- well plates to an OD_600_ of 0.2, as determined by a Biotek Synergy HT microplate reader. A total of eight 1:10 serial dilutions were made with sterile ddH_2_O, and 3 μL of cells were plated onto YPD, 50, 100 and 200 mM hydroxyurea, and 150, 250 μM 5-fluorouracil. Plates were grown for three days at 30°C and imaged. Quantitative efficiency of plating assays were performed by taking dilutions of the above yeast strains and spreading 100 μL cells on YEPD, 50 mM HU, and 150 μM 5-FU media. Plates were grown for three days at 30°C and colonies were counted on each plate. Three biological replicates of each dilution and media type were analyzed.

**Table 1.**
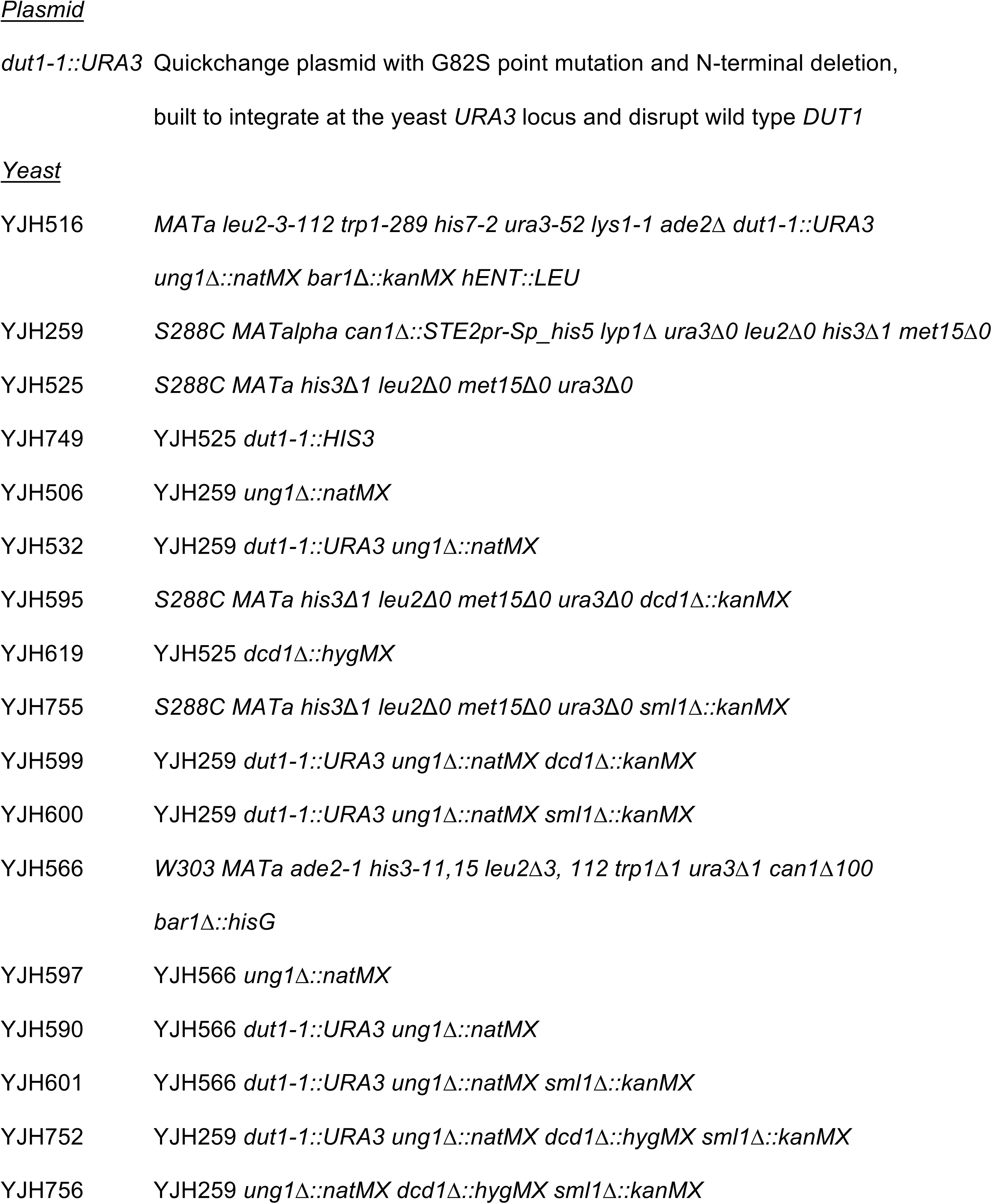
Yeast strains and plasmids.

### Excision-seq library methods

*S. cerevisiae* (YJH532, 599, 600, and 601, **Table 1**) were used to prepare post-digestion Excision-seq libraries. These libraries used adaptors JH0804 and JH1139 (**Table 2**) and were PCR amplified with primers JH1140 and JH1159, 1162, or 1163 (**Table 2**).

**Table 2.**
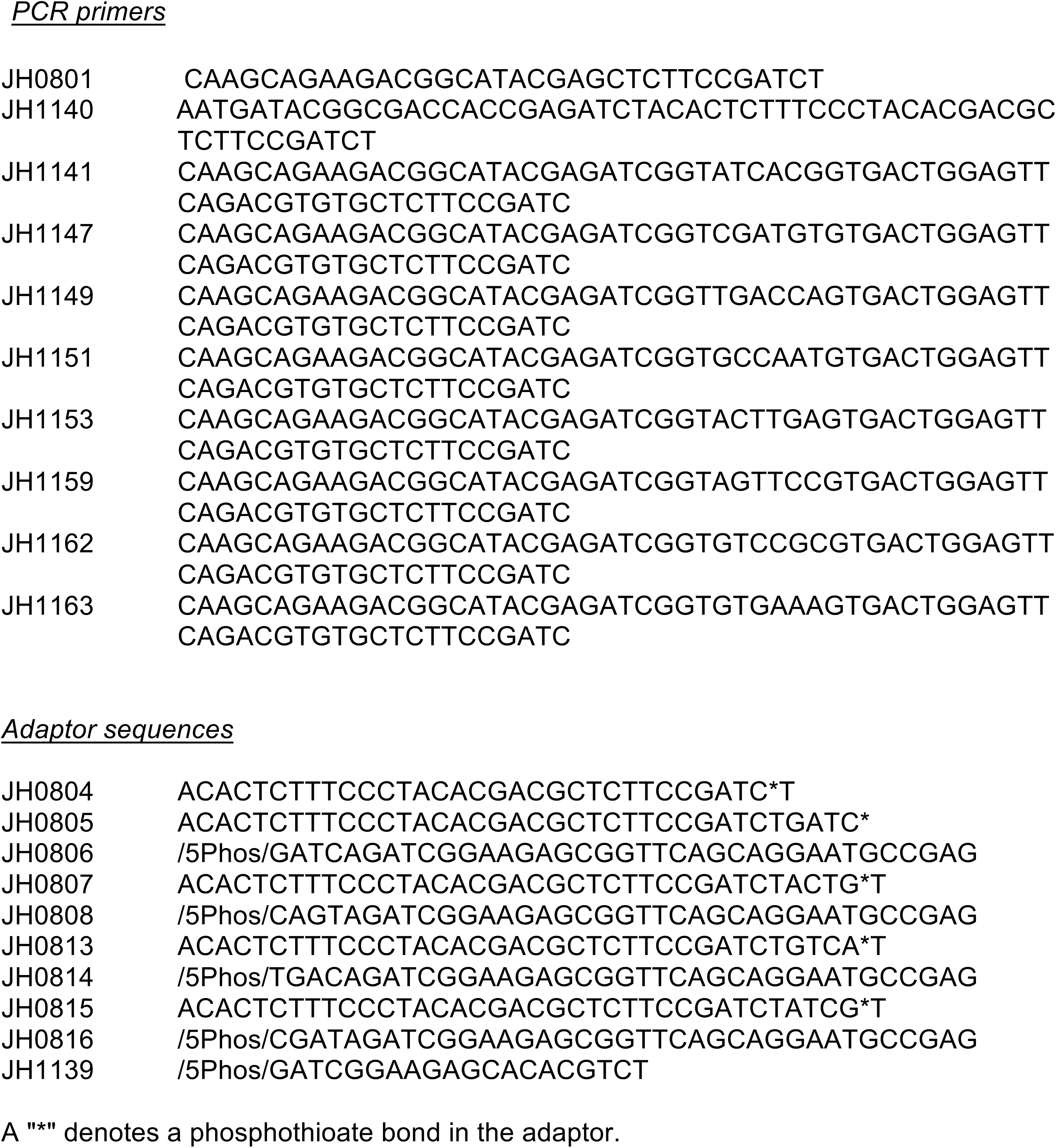
Oligonucleotides.

Detailed methods for constructing Excision-seq libraries are available in Bryan et al. 2014. Briefly, in post-digestion Excision-seq, libraries of mechanically sheared genomic DNA were treated with excision repair enzymes to destroy strands containing the modified base, preventing their PCR amplification. Libraries were sequenced on Illumina MiSeq or HiSeq 2000 platforms using standard protocols. Sequencing data are available at NCBI GEO under accession number GSE73364.

### Analysis of Excision-seq data

Sequences were analyzed by alignment to a reference genome (*sacCer1*) using Bowtie (Langmead et al. 2009) and SAMtools (Li et al. 2009), processed to bedGraph format using BEDTools (Quinlan *et al.* 2010), and visualized in the UCSC Genome Browser (Karolchik *et al.* 2001). Coverage at each position was normalized by the number of reads aligned in the library (*i.e.,* reads per million, RPM). Using this method, the level of coverage at a specific site or region in the genome represents the relative quantity of modified base at that position. Software and pipelines used to analyze data are available on GitHub (https://github.com/hesselberthlab/modmap).

